# Music elicits different gene expression responses in the buccal cavity of age-related cognitive disorders patients and healthy controls

**DOI:** 10.1101/2024.05.29.596389

**Authors:** Alberto Gómez-Carballa, Laura Navarro, Nour El Zahraa Mallah, Xabier Bello, Sara Pischedda, Sandra Viz-Lasheras, María José Currás, Isabel Ferreirós-Vidal, Narmeen Mallah, Julián Montoto-Louzao, Alba Camino-Mera, Lúa Castelo-Martínez, Sara Rey-Vázquez, Lorenzo Redondo, Ana Dacosta-Urbieta, Irene Rivero-Calle, Carmen Rodriguez- Tenreiro, Federico Martinón-Torres, Antonio Salas, Sensogenomics Working Group

## Abstract

Recent evidence suggests that external stimuli can shape transcriptomes (sensogenomics). Specifically, the analysis of capillary blood samples has shown that musical stimuli can modulate gene expression patterns, in healthy individuals but also in those with age-related cognitive disorders (ACD), based on. Here, we present groundbreaking evidence indicating that brief exposure to music can also impact the salivary transcriptome in both healthy donors and ACD patients. Our findings reveal that music has a more pronounced effect on patients compared to controls, inducing global gene expression changes towards upregulation in ACD patients but downregulation in controls. The most significantly dysregulated genes in ACD patients include *LGALS3* (downregulated) and *CXCL8* (upregulated), whereas in controls, *THOP1* was the top significant gene (downregulated). These genes play important roles in normal brain functions and are also altered in neurodegenerative conditions. Weighted Gene Co-expression Network Analysis reveals relevant and significant modules, both positive and negative correlated with music, implicated in neurodegenerative (e.g. autophagy) and immunological processes (e.g. IL-1, MHC). This sheds light on the complex interplay between music and molecular responses in the human body. This study underscores the potential of musical stimuli to influence gene expression patterns outside of systemic circulation, paving the way for further exploration of music’s therapeutic effects.

**Sensogenomics Working Group:** Antonio Salas Ellacuriaga – PI; Federico Martinón-Torres – PI; Laura Navarro Ramón – Coordinator

*GenPoB/GenVip - Instituto de Investigación Sanitaria (IDIS) (alphabetic order):* Alba Camino Mera, Albert Padín Villar, Alberto Gómez Carballa, Alejandro Pérez López, Alicia Carballal Fernández, Ana Cotovad Bellas, Ana Isabel Dacosta Urbieta, Narmeen Mallah, Ana María Pastoriza Mourelle, Ana María Senín Ferreiro, Andrés Muy Pérez, Antía Rivas Oural, Antonio Justicia Grande, Antonio Piñeiro García, Anxela Cristina Delgado García, Belén Mosquera Pérez, Blanca Díaz Esteban, Carlos Durán Suárez, Carmen Curros Novo, Carmen Gómez Vieites, Carmen Rodríguez-Tenreiro Sánchez, Celia Varela Pájaro, Claudia Navarro Gonzalo, Cristina Serén Trasorras, Cristina Talavero González, Einés Monteagudo Vilavedra, Estefanía Rey Campos, Esther Montero Campos, Fernando Álvez González, Fernando Caamaño Viñas, Francisco García Iglesias, Gloria Viz Rodríguez, Hugo Alberto Tovar Velasco, Irene Álvarez Rodríguez, Irene García Zuazola, Irene Rivero Calle, Iria Afonso Carrasco, Isabel Ferreirós Vidal, Isabel Lista García, Isabel Rego Lijo, Iván Prieto Gómez, Iván Quintana Cepedal, Jacobo Pardo Seco, Jesús Eirís Puñal, José Gómez Rial, José Manuel Fernández García, José María Martinón Martínez, Julia Cela Mosquera, Julia García Currás, Julián Montoto Louzao, Lara Martínez Martínez, Laura Navarro Marrón, Lidia Piñeiro Rodríguez, Lorenzo Redondo Collazo, Lúa Castelo Martínez, Lucía Company Arciniegas, Luis Crego Rodríguez, Luisa García Vicente, Manuel Vázquez Donsión, María Dolores Martínez García, María Elena Gamborino Caramés, María Elena Sobrino Fernández, María José Currás Tuala, María Martínez Leis, María Soledad Vilas Iglesias, María Sol Rodriguez Calvo, María Teresa Autran García, Marina Casas Pérez, Marta Aldonza Torres, Marta Bouzón Alejandro, Marta Lendoiro Fuentes, Miriam Ben García, Miriam Cebey López, Montserrat López Franco, Nour El Zahraa Mallah, Narmeen Mallah, Natalia García Sánchez, Natalia Vieito Perez, Patricia Regueiro Casuso, Ricardo Suárez Camacho, Rita García Fernández, Rita Varela Estévez, Rosaura Picáns Leis, Ruth Barral Arca, Sandra Carnota Antonio, Sandra Viz Lasheras, Sara Pischedda, Sara Rey Vázquez, Sonia Marcos Alonso, Sonia Serén Fernández, Susana Rey García, Vanesa Álvarez Iglesias, Victoria Redondo Cervantes, Vanesa Álvarez Iglesias, Wiktor Dominik Nowak, Xabier Bello Paderne, Xabier Mazaira López

*Nursing volunteers (alphabetic order):* Alejandra Fernández Méndez, Ana Isabel Abadín Campaña, Ana María León Caamaño, Ana María Buide Illobre, Ángeles Mera Cores, Carmen Nieves Vastro, Carolina Suarez Crego, Concepción Rey Iglesias, Cristina Candal Regueira, Dolores Barreiro Puente, Elvira Rodríguez Rodríguez, Eugenia González Budiño, Eva Rey Álvarez, Fernando Rodríguez Gerpe, Gemma Albela Silva, Isabel Castro Pérez, Isabel Domínguez Ríos, José Ángel Fernández de la Iglesia, José Cruces Vázquez, José Luis Cambeiro Quintela, José Ramón Magariños Iglesias, Julia Rey Brandariz, Julio Abel Fernández López, Luisa García Vicente, Manuel González Lito, Manuel González Lijó, Manuela Pérez Rivas, Margarita Turnes Paredes, María Aurora Méndez López, María Begoña Tomé Arufe, María Campos Torres, María del Carmen Baloira Nogueira, María del Carmen García juan, María Esther Moricosa García, María Luz Chao Jarel, María Martínez Leis, María Mercedes Jiménez Santos, María Salomé Buide Illobre, María Victoria López Pereira, Mercedes Jorge González, Mercedes Isolina Rodríguez Rodríguez, Miren Payo Puente, Natalia Carter Domínguez, Olga María Reyes González, Pilar Mera Rodríguez, Purificación Sebio Brandariz, Salomé Quintáns lago, Yolanda Rodríguez Taboada, María Pereira Grau.

*Other volunteers (alphabetic order):* Alba Arias Gómez, Alejandro Moreno Díaz, Ana Arca Marán, Astro González Guirado, Brais García Iglesias, Carlos Sánchez Rubín, Carmen Otero de Andrés, Clara Pérez Errazquin Barrera, Claudia Rey Posse, Cristina Rojas García, Eduardo Xavier Giménez Bargiela, Elena Gloria Morales García, Fabio Izquierdo García Escribano, Gabriel Guisande García, Jaime López Martín, Lara Pais Ramiro, Lucía Rico Montero, Luís Estévez Martínez, Manuel Estévez Casal, María Aránzazu Palomino Caño, María Rubio Valdés, Marisol Nogales Benítez, Miryam Tilve Pérez, Nuria Villar Muiños, Pablo Del Cerro Rodríguez, Pablo Pozuelo Martínez Cardeñoso, Salma Ouahabi El Ouahabi, Santiago Vázquez Calvache

## Introduction

Little is known about how musical stimuli impacts on our genetic expression. Navarro et al. (1) have recently underscored the importance of deeper exploration into this still poorly understood field of biological science (sensogenomics (1-3)), taking advantage of new technologies emerging in the ‘-omic’ sciences, including genomics and transcriptomics. There have been only a few attempts to understand the gene expression mechanisms activated during music stimulation; the initial ones were carried out on healthy controls and professional musicians, with results indicating a few genes differentially expressed after exposure to classical music stimuli (4, 5). Recently, the exploratory study by Navarro et al. (3) provided suggestive evidence of the potential impact of music in the context of Alzheimer’s disease (AD) and reviewed the current evidence overall, concurring on the beneficial effect of music on neurodegenerative diseases. To the best of our knowledge, our most recent study, Gómez-Carballa et al (2), is the only attempt to date to examine the impact of music in a disease context, specifically in capillary blood samples collected from age-related cognitive disorder (ACD) patients. This study (2) found an increased effect of music in ACD patients compared to healthy controls, but most interestingly, it revealed that brief musical stimuli can modify the way patients express genes typically altered in this condition, but in the opposite direction.

Building upon previous findings, the present study represents the first attempt to analyze the impact of music in ACD patients, but this time exploring salivary instead of blood transcriptomes. The interest in analyzing saliva stems from the growing importance of this non-invasive biological source in biomedical studies, with several salivary biomarkers being explored as proxies for diagnosing and monitoring brain health, stress, mental disorders, and neurological diseases (6-10). Saliva is primarily produced and secreted by the parotid, submandibular, and sublingual salivary glands and regulated by the autonomous nervous system (11). These major salivary glands are innervated by both sympathetic and parasympathetic nerves, with compact fibers encircled by Schwann cells (12). The submandibular and sublingual glands are responsible for most unstimulated saliva production, whereas most of parotid gland saliva secretion occurs in response to stimuli. Previous studies have reported the presence of neurotransmitters in salivary glands extracted from mice and rats (13, 14). Parasympathetic stimulation promotes the release of the neurotransmitter acetylcholine whereas sympathetic activation releases noradrenaline stimulating the secretion of proteins (15, 16). Given this connection between the salivary glands and the nervous system, the existence of a bidirectional oral-brain axis has been suggested (17), through which an inflammatory response in the oral cavity may impact brain homeostasis and vice-versa. Since there is growing evidence indicating an impact of music on blood transcriptomes as well as its beneficial effects on many disease conditions, it seems imperative to explore if musical stimuli have also the potential to regulate salivary transcriptomes.

Here, we propose exploring gene expression patterns in saliva obtained from ACD patients and healthy controls before and after brief musical stimuli. The samples analyzed in this study partially overlap with those used in Gómez-Carballa et al. (2); and all of them were collected during the same experimental concerts. Therefore, this scenario offers a unique opportunity to evaluate the potential of saliva analysis in capturing expression changes triggered by music, and to undertake a comparative analysis with the capillary blood signatures reported in our previous study (2).

## Methods

### Participants and *n*Counter assay

Written informed consent was obtained from all the participants in the present study. The Ethics Committee of Xunta de Galicia approved the present project (Registration code: 2020/021), and the study was conducted in accordance to the guidelines of the Helsinki Declaration. We have followed the same experimental procedures described in Gómez-Carballa et al. (2) and within the framework of the Sensogenomics project (http://sensogenomics.com). Briefly, we collected saliva samples at two timepoints: before and after an experimental concert of classical music; see Figure 1 of (2) for a schematic representation of the sampling and analysis procedures. Saliva samples were collected in *Oragene DNA* devices (ORE-100; DNAgenotek), comprising 10 ACD patients (aged 84-92 years old; mean 84) and 14 healthy donors (aged 18-88; mean 57). Nearly all the ACD patients (9/10; 90%) and more than half of the controls (8/14; 57%) from the present study overlap with those included in the capillary blood experiment (2).

**Figure 1.**
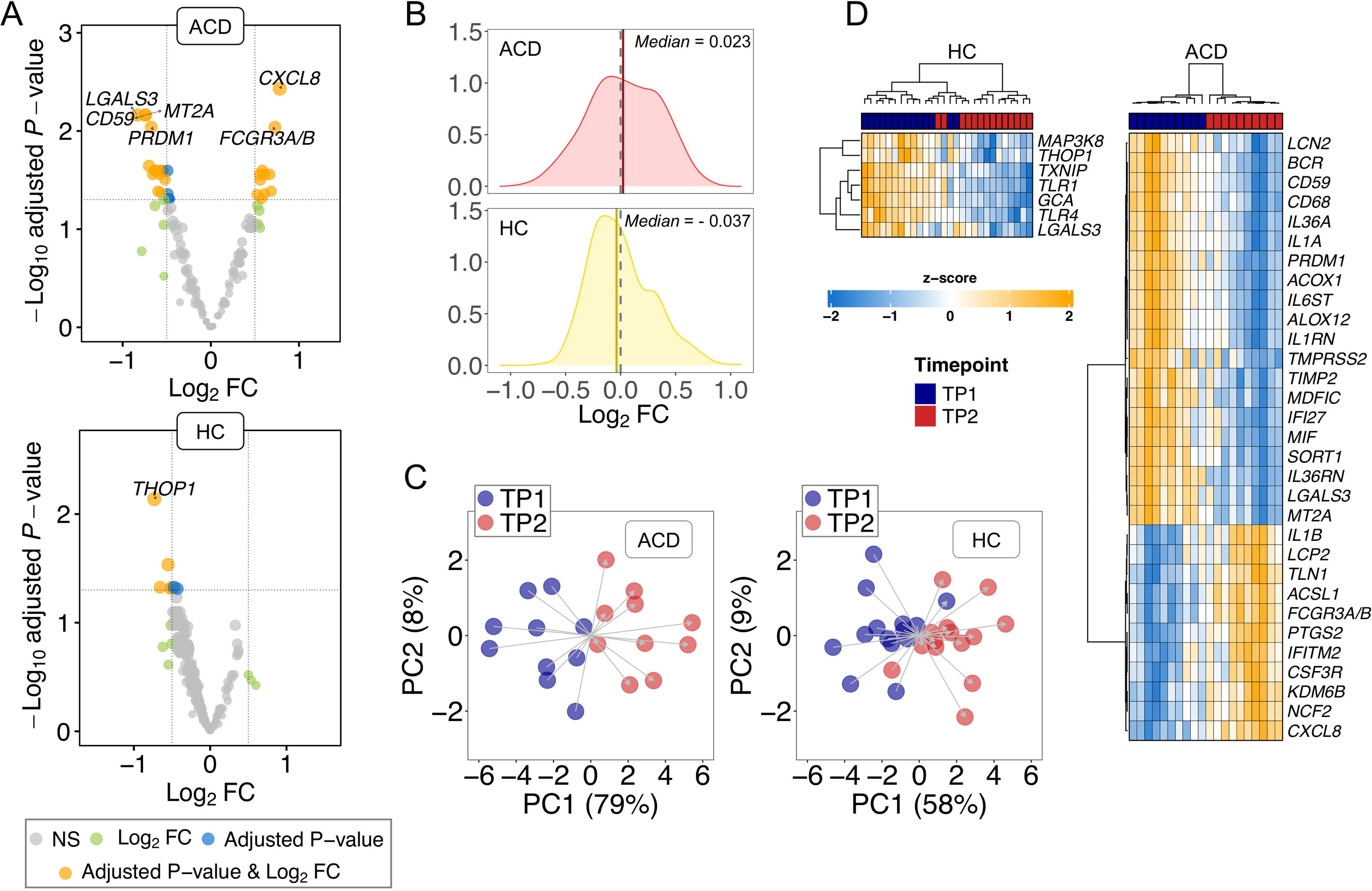
A) Volcano plot showing the DEGs between TP1 and TP2 in both ADC and controls donors (HC). Names of DEGs with |log_2_FC| value > 0.5 and adjusted *P*-value < 0.01 are displayed. B) Density plots of log_2_FC; dashed lines indicate Log_2_FC = 0; vertical solid lines indicate median values; C) Principal components analysis (PCA) using DEGs with *P*-value < 0.05. D) Heatmap and cluster analysis of the DEGs (adjusted *P*-value < 0.05) in ADC and HC.

RNA from saliva was isolated using 500µl of sample and the RNeasy microkit (Qiagen). We slightly modified the protocol provided by the extraction kit as recommended by the Oragene tubes supplier. RNA concentration step and an additional DNase treatment were undertaken using an RNA clean & concentrator kit (Zymo Research). RNA amount and integrity were checked using TapeStation 4200 (Agilent), and DV200 values were calculated to ensure that >50% of the RNA fragments were above 200nt and to estimate the optimal sample input.

Gene expression was evaluated through the *nCounter MAX* (NanoString Technologies) and the *n*Counter Host Response Panel, which includes 785 genes. We opted for NanoString over other methods such as RNAseq due to the intrinsic difficulties associated to sequencing endogenous RNA from saliva samples. Bacteria are naturally present in human saliva, making it challenging to analyze only the human component of the salivary transcriptome (18). We followed standard protocols; including 12× RNA hybridization with 5 μl of RNA as input, and hybridization time of 18 hours for all samples. We also mixed controls and ACD patient samples in the same runs to avoid technical sample bias and batch effects. After filtering out genes expressing below the background (maximum expression value < background), we detected a total of 566 and 672 genes (out of the total 785 in the NanoString panel) in ACD patients and healthy controls, respectively. In addition, 553 out of 566 genes in ACD and 648 out of 672 genes in healthy controls overlap with those detected in capillary blood samples from our previous study (2).

### Statistical analysis

First, we carried out a quality control (QC) of the raw expression data checking some technical parameters following manufacturer recommendations to verify the absence of technical problems. Samples that did not pass technical QC, or with low number of genes detected, were excluded for downstream analysis.

Genes with counts below the background (defined as the mean + 2 standard deviations [SD] of the negative control spikes in the code set, disregarding negative control C, which typically yields a higher number of counts) were excluded from both normalization and the differential expression analysis.

Data normalization was performed through an iterative strategy that combines both *DESeq2* (19) and *RUVSeq* (20) packages as described in (21). Control reference genes for data normalization were detected by selecting invariable genes (*P*-value > 0.1, BaseMean > 100 and |log_2_FC| < 0.2) after a naïve differential expression analysis between TP1 And TP2 in both ACD and healthy controls separately. Genes expressed below the background were removed. We used a paired-sampling design to carry out analysis of transcriptome differences before the musical stimuli (Pretest; time-point one or TP1) and after the musical stimuli (Posttest; TP2). Additionally, we evaluated if DEGs detected in saliva are also altered in AD/MCI patients due to the condition by contrasting DEGs between TP1 and TP2 against DEGs that are altered in AD and mild cognitive impairment (MCI) patients. For this purpose, we downloaded from GEO (gene expression omnibus) microarray blood gene expression data from three independent microarray datasets analyzing MCI and AD patients and healthy controls, as done previously (2). Data were processed and merged as explained in (2, 3).

We used the Weighted Gene Co-expression Network Analysis (WGCNA) R package (22) to investigate clusters of co-expressed genes potentially correlated to the musical stimuli in ACD patients and the healthy control groups separately. Normalized and corrected gene expression data, adjusted for patient-to-patient variability, served as input to construct a signed weighted correlation network. Following the package developers’ recommendations and considering the number of samples per group, we chose a soft-thresholding power of 18. We computed the Topological Overlap Matrix (TOM) and the corresponding dissimilarity (1–TOM) values. We set a minimum module size of 30, and a dendrogram cut height threshold of 0.2 for module merging. Initially labeled by colors, the detected modules of co-expressed genes were later renamed using the name of the genes showing the highest connectivity within each module (hub genes). We identified modules of interest significantly associated with the musical stimuli by correlating module eigengenes with the time point (TP1 and TP2) data; and measuring gene significance (GS), a value that quantifies the biological significance of genes in modules. For each gene, Module Membership (MM) quantifies its intramodular connectivity within the module. Multiple test adjustment was carried out using the FDR method by Benjamini-Hochberg (23).

Functional analysis of significantly correlated modules was carried out through an over-representation analysis with the *Clusterprofiler* R package (24). The biological processes from the Gene Ontology (GO) and Reactome were used as reference databases for the analysis. The pool of genes included in the *n*Counter NanoString Host Response gene expression panel was employed as the gene universe for statistical calculations. To facilitate the interpretation of the results, redundant terms (similarity > 0.7) were detected and removed after calculating the terms similarity matrix.

Different R packages were used to generate volcano plots (*EnhancedVolcano* (25)) and heatmaps (*ComplexHeatmap* (26)). Statistical significance was assessed using the Wilcoxon test.

Statistical analyses were performed using R version 4.2.2. (27).

## Results

### Differentially expression in saliva in response to music

To assess the impact of the musical stimuli on the transcriptomes of donors, we first conducted a paired TP1 *vs*. TP2 transcriptome analysis for the two groups of donors separately.

First, we observed a higher number of DEGs in ACD patients compared to controls. Specifically, we detected 31 (adjusted *P*-value < 0.05) DEGs in the ACD group out of a total of 566 detected genes. In contrast, we found 7 adjusted DEGs in healthy controls out of a total of 672 detected genes. These different proportions (31/566 *vs*. 7/672) were statistically significant under a two-sample proportion test (*P*-value = 1×10^-05^); **Table S1, Table S2, Figure 1A**.

Secondly, the musical stimuli drive the transcriptome of ACD patients towards upregulation when compared to controls. Notably, there are more upregulated adjusted DEGs in ACD donors (11/31 = 0.35) compared to the lower proportion observed in the healthy controls (0/7). Moreover, upregulation appears to be the predominant overall response to music in the transcriptome of ACD patients (median log_2_FC = 0.032 of non-adjusted DEGs with *P*-value < 0.05), whereas downregulation predominates in the transcriptome of healthy donors (median log_2_FC = –0.037 of DEGs with non-adjusted *P*-value < 0.05); **Figure 1B**. While these figures are inadequate for a two-sample proportion test (as it is an incorrect approximation to a Chi-square), we conducted the test using the observed non-adjusted DEGs, with proportions 52/97 = 0.54 in ACD patients *vs*. 45/128 = 0.35 in healthy controls; these different proportions were statistically significant with *P*-value = 0.009.

The transcriptome profiles of non-adjusted DEGs reveal a segregation of samples into their two timepoints (TP1 and TP2) in ACD patients, as evidenced by a PCA. This differentiation is particularly noticeable in Principal Component 1 (PC1), which account for most of the variation (79%); PC2 contributes minimally to this primary PC1 clustering, representing only 8% of the variation; **Figure 1C**. However, the differentiation between TP1 and TP2 is less pronounced in the healthy cohort, with PC1 and PC2 accounting for only 58% and 9% of the variation, respectively; **Figure 1C**. A heatmap of DEGs (adjusted *P*-value < 0.05) in ACD clearly demonstrates a clear distinction between TP1 and TP2 in ACD patients. Despite the limited number of DEGs observed in healthy donors (*n* = 7), the heatmap efficiently separates most of the transcriptomes into the two timepoints; **Figure 1D**.

The top downregulated gene in ACD was *LGALS3* (log_2_FC = –0.83; adjusted *P*-value = 0.007) whereas the DEG showing the lowest adjusted *P*-value was *CXCL8* (log_2_FC = 0.78; adjusted *P*-value = 0.004). In control samples, the top DEG, namely *THOP1*, was downregulated in TP2 with respect to TP1 (log_2_FC = – 0.73; adjusted *P*-value = 0.007); **Table S1**.

Using a pathways enrichment approach and DEGs (adjusted *P*-value < 0.05) a single GO significant term was detected in ACD patients: “unsaturated fatty acid metabolic process” (adjusted *P*-value = 0.03), involving the DEGs *IL1B*, *PTGS2*, *ACOX1*, *MIF* and *ALOX12*. Nonsignificant pathway resulted from the analysis in control donors, most likely due to the low number of DEGs detected.

### Comparative transcriptomic response to music in capillary blood and saliva

We observed few similarities between transcriptomic response in the capillary blood and saliva of donors exposed to musical stimuli (comparing the values mentioned above with results reported in (2); see also **Table S2**); namely: *i*) There is a statistically significant higher number of DEGs in ACD patients compared to controls, *ii*) Global upregulation is the predominant reaction to music in the transcriptome of ACD patients, while downregulation predominates in the transcriptome of healthy donors, and *iii*) The transcriptome profile of DEGs shows a clear distinct differentiation between TP1 and TP2 in both healthy controls and, even more marked, in ACD patients.

However, there are also a few differences between the transcriptomes in saliva and capillary blood of patients and controls that are worth highlighting.

Firstly, the most notable finding is that the proportion of adjusted DEGs is substantially higher in saliva than in blood (**Table S2)**, for both ACD patients and healthy controls. When referred to the total number of transcripts captured by the different techniques (NanoString in saliva and RNAseq in capillary blood (2)), the proportions are as follows: *i*) in ACD patients: 31/566 = 0.05 [saliva] *vs*. 328/36155 = 0.01 [capillary blood]; and *ii*) in healthy controls: 7/672 = 0.01 [saliva] *vs*. 1/35865 = 0.00 [capillary blood]. For these comparisons, the two-sample proportion test is highly significant, *P*-value < 2×10^-16^; **Table S2**. In addition, to mitigate potential bias arising from the different techniques employed to generate salivary and capillary blood transcriptomes, we can consider only the common genes in both studies (553 in ACD patients and 648 in healthy controls; see Material and Methods). Although the proportions are more attenuated, they remain consistent: *i*) in ACD patients: 30/553 = 0.05 [saliva] *vs*. 21/553 = 0.04 [capillary blood], and *ii*) in healthy controls: 7/648 = 0.01[saliva] *vs*. 0/648 = 0.00 [capillary blood]. This difference is statistically significant in healthy controls (*P*-value = 0.02); but it is highly significant in both groups when computing proportions using non-adjusted DEGs; **Table S2**.

Secondly, we observed a significant proportion of genes that express in different directions in capillary blood and saliva. Specifically, there are 24 common (non-adjusted) DEGs in the two tissues for ACD patients, and 7 in healthy controls. Among these, 11/24 = 0.46 [ACD patients], and 2/7 = 0.29 [controls] were found to be negatively correlated (**Figure 2**). Additionally, there are only two DEGs (adjusted *P*-value < 0.05) in ACD patients (none in controls), namely *KDM6B* and *TIMP2,* that overlap between saliva and blood transcriptomes. While *TIMP2* expressed similarly in saliva and capillary blood, *KDM6B* is upregulated in saliva but downregulated in blood; **Figure 2**.

**Figure 2.**
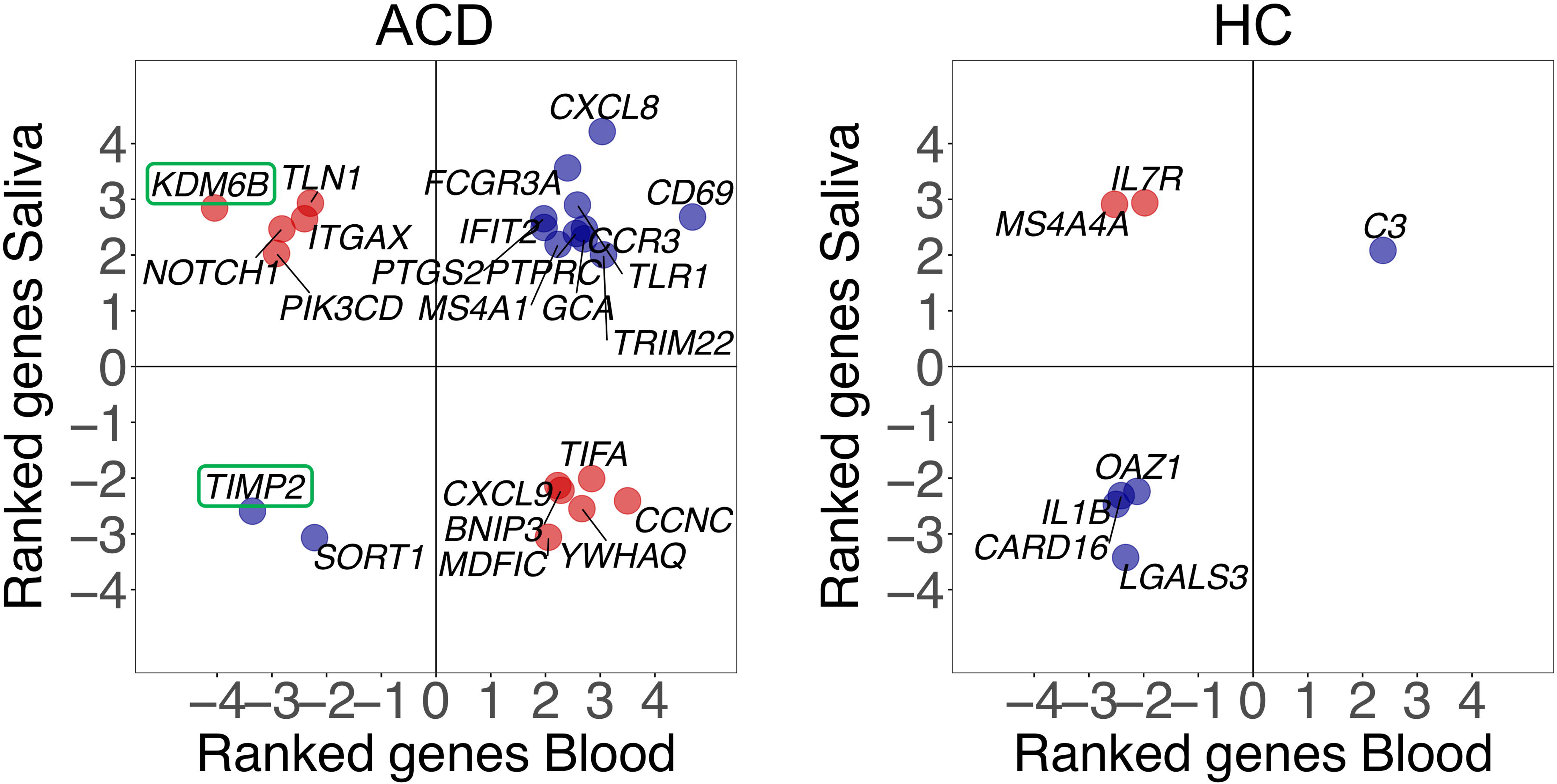
Correlation between gene expression changes observed in capillary blood and saliva samples after musical stimulation in ACD and control donors (HC). Blue dots indicate positive correlation whereas blue dots indicate negative correlation between tissues. Only common DEGs (*P*-value < 0.05) in both saliva and capillary blood samples are being displayed. Ranked values refer to the value of the test statistic for a gene obtained from Deseq2.

**Figure 2.**
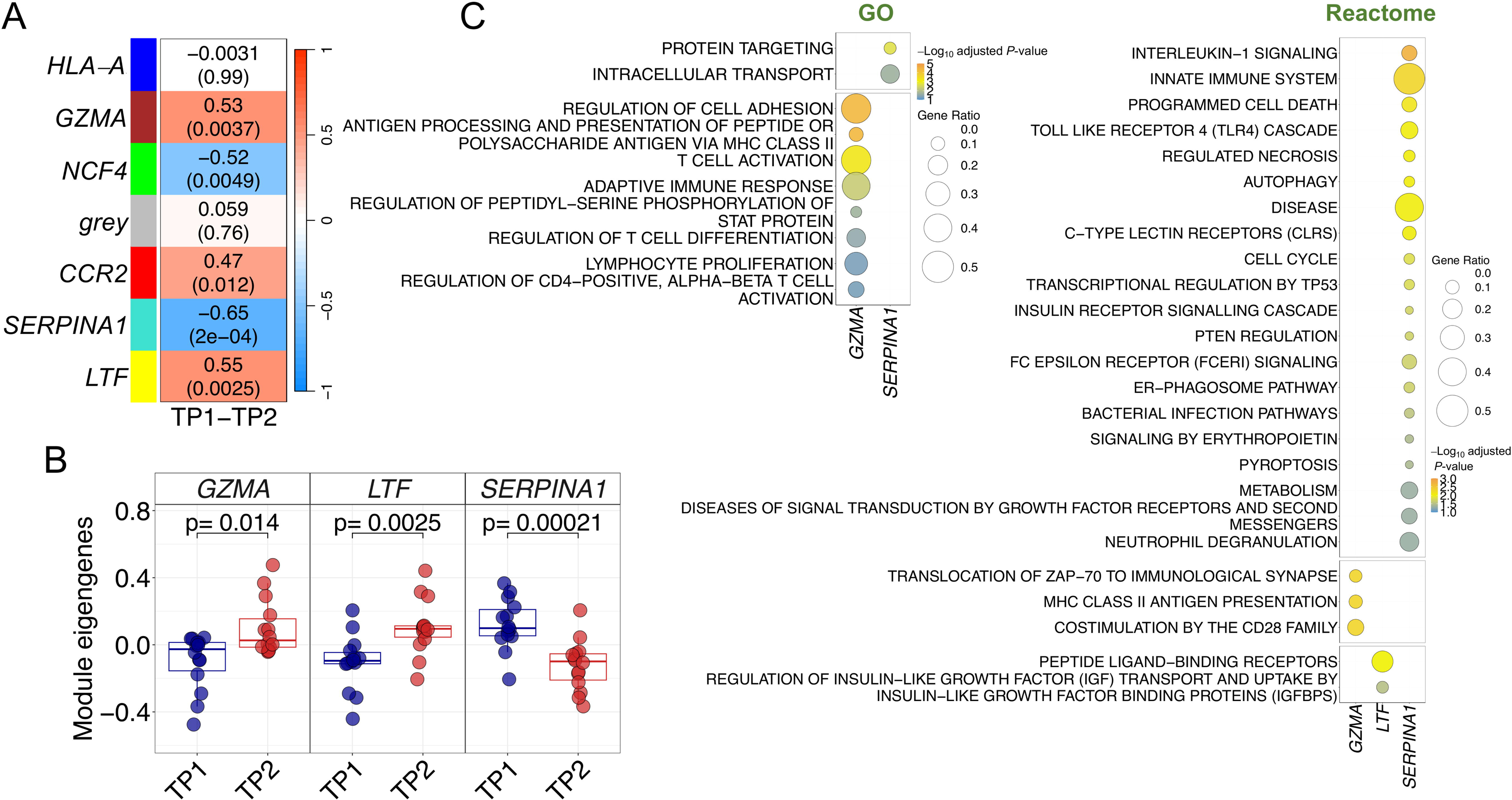
Co-expression analysis in control donors. (A) Correlation and *P*-values values heatmap obtained from the co-expression analysis. Upper value corresponds to the correlation with musical stimuli and *P*-values are shown in brackets (B) Boxplots of samples eigengenes values between TP1 and TP2 from *GZMA*, *LTF* and *SERPINA1* modules. (C) Pathways enrichment analysis results using GO and Reactome databases.

### Music-related DEGs in saliva from ACD patients compared to DEGs in AD/MCI condition

To investigate if some of the genes affected by musical stimuli in ACD were also dysregulated in AD/MCI patients due to their condition, we contrasted the DEGs detected in ACD after the musical stimuli with DEGs resulted from comparing transcriptomes from AD/MCI patients and healthy controls. We observed that some genes targeted by music were also affected in both neurodegenerative conditions. However, music appeared to impact more significantly on genes dysregulated in MCI (*n* = 13) than in genes dysregulated in AD (*n* = 8); **Figure S1**. There were few genes that showed a negative correlation in these contrasts, more in MCI (*n* = 7) than in AD (*n* = 3); suggesting that music has a compensatory effect for those altered in the two disease conditions.

Interestingly, the only significant genes altered by music in both capillary blood and saliva (see above) were also differentially expressed in AD and MCI, namely *KDM6B* and *TIMP2* (**Figure S1)**. Both genes showed a significant higher expression in MCI and AD compared to healthy controls. However, music induced opposite expression changes in saliva for the *KDM6B* (over-expression as in MCI/AD) and *TIMP2* (under-expression) genes.

### Co-expression modules in response to the musical stimuli in ACD

The WGCNA analysis generated six modules of co-expressed genes from the ACD expression data (**Figure 3A; Figure S2A** and **S2B; Table S3**). Correlation of the modules eigengenes with TP1 and TP2 revealed four modules significantly correlated with the expression changes produced by the music stimuli (**Figure 3A**). Three of them were positively correlated: *PIK3CD* ([blue] *P*-value = 0.004), *NOTCH1* ([brown] *P*-value = 5×10^-06^), and *CXCR1* ([green] *P*-value = 0.002); whereas the module *CD59* ([turquoise] *P*-value = 7×10^-05^) showed a negative correlation. *NOTCH1* and *CD59* modules showed the highest correlation with the musical stimuli (*R* = 0.83 and *R* = –0.77, respectively).

**Figure 3.**
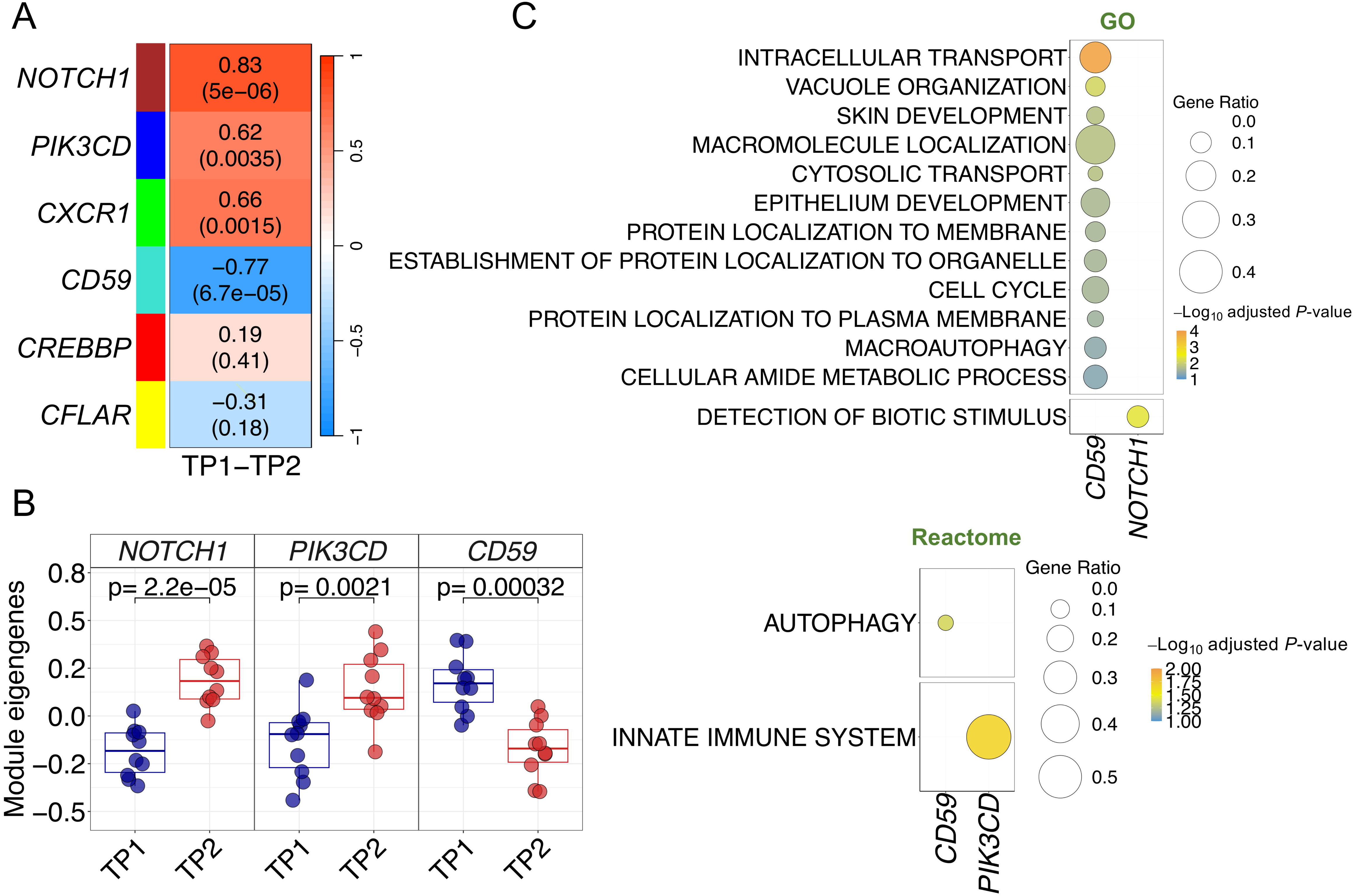
Co-expression analysis in ACD patients. (A) Correlation and *P*-values values heatmap obtained from the co-expression analysis. Upper value corresponds to the correlation with musical stimuli and *P*-values are shown in brackets (B) Boxplots of samples eigengenes values between TP1 and TP2 from *NOTCH1*, *CD59* and *PIK3CD* modules. (C) Pathways enrichment analysis results using GO and Reactome databases.

The highly significant correlation values of these four modules suggest a functional role of these gene sets in the buccal molecular response to music in ACD patients. This functional relevance is likewise visible by examining the global correlation between gene MM and gene-trait correlation values for the individual genes within each module, indicating that genes with higher trait-correlation values are also important functional drivers of the modules (high MM values) (*R* = 0.73 and *P*-value = 9×10^-34^ for *CD59* module; *R* = 0.74 and *P*-value = 3×10^-14^ for *NOTCH1* module); **Figure S2C**. Modules *NOTCH1*, *CD59* and *PIK3CD* are the ones showing the most significant expression changes between TP1 and TP2 (**Figure S2D**); this is particularly clear when examining their module eigengene values (**Figure 3B**).

Pathways enrichment analysis of the significant modules identified relevant biological routes involved in the musical stimuli for three of the modules; *CD59*, *NOTCH1* and *PIK3CD* (**Table S4; Figure 3C**). The most relevant pathways detected for the downregulated *CD59* module were related to intracellular protein transport and localization (in GO) and, most remarkable, autophagic machinery/vacuole organization (in GO and Reactome); **Figure 3C**. Genes from the upregulated *NOTCH1* and *PIK3CD* modules were involved in the detection of biotic stimulus (*NOTCH1* in GO) and innate immune system (*PIK3CD* in Reactome); **Figure 3C**.

### Co-expression modules in response to the musical stimuli in controls

The co-expression module analysis in control donors also detected significant modules altered in response to music, but the correlations were generally lower than those reported for the ACD patients (**Figure 4A; Figure S3A** and **S3B; Table S3**). Three modules yielded positive correlation values (*LTF* - [yellow] *P*-value = 0.002, *GZMA* - [brown] *P*-value = 0.004, *CCR2* - [red] *P*-value = 0.01), indicating upregulation after receiving musical stimuli (in TP2), whereas two of them showed negative correlation values (*SERPINA1* - [turquoise] *P*-value = 0.0002, *NCF4* - [green] *P*-value = 0.005), indicating downregulation after listening to music; all of them survived adjustment for multiple test (**Table S3**). Among the topmost significant modules (*SERPINA1*, *GZMA* and *LTF*), the most significant one was also the most negatively correlated with musical stimulation (*SERPINA1*; *R* = – 0.64), and the module displaying the most extreme differences in eigengenes values between both timepoints (*P*-value = 0.0002); **Figure 4B**. Although the overall gene expression profiles of the top three modules clustered reasonably well the samples into TP1 and TP2 (**Figure S3C**), this separation by TP was not as optimal as in the case of the modules from ACD patients. The importance of the *SERPINA1* module and its core genes in the gene expression response of control donors can also be noted in the high correlation and low significance values observed when contrasting gene MMs with GSs individual values (*R* = 0.70; *P*-value = 1×10^-34^) in comparison to the values obtained for the other modules (**Figure S3D**).

Several pathways were found to be related to *GZMA*, *SERPINA1* and *LTF* modules after a functional assessment of each of the significantly correlated modules (**Figure 4C; Table S4**). The *GZMA* module showed a strong involvement of the adaptative immune system. Thus, most significant pathways can be condensed into three sets of adaptive immune-related processes: antigen processing and presentation *via* MHC class II, cell adhesion, and T-cell metabolism routes (activation, proliferation, regulation, differentiation). Consistently, enrichment analysis with Reactome database yielded equivalent results, with MHC class II antigen presentation as the top significant pathway, and other pathways with a key role modulating T-cell activity (CD28, PD-1 signalling) or triggering downstream cascades after T-cell receptor activation (ZAP-70); (**Figure 4C; Table S3**). GO enrichment analysis of *SERPINA1* module resulted in intracellular protein transport and related terms significantly associated with this module. However, Reactome enrichment analysis found several significant pathways engaged in the innate immune system, locating at the top different Toll-Like Receptors (TLR) cascades and related pathways, such as IL-1 signalling, MyD88:MAL (TIRAP) cascade or TRAF6 mediated induction of NFkB and MAP kinases. Most noticeable, autophagy pathways were also significantly associated with the *SERPINA1* module (autophagy, macro-autophagy and ER-Phagosome pathways) suggesting functional similarities with the negatively correlated module *CD59* detected in ACD patients (see above). Processes associated with the *LTF* module were only detected in Reactome; related to the GPCR signal transduction events, and more specifically to the sub-family A/1 (Rhodopsin-like) receptors.

## Discussion

Musical stimulation is a multifaceted cognitive phenomenon that intricately engages various brain regions, eliciting a spectrum of cognitive, emotional, and physiological responses. Elucidating the molecular changes triggered by music can help to understanding its effect on brain function and mental health in the general population, as well as its therapeutic potential in the context of various neurological and psychiatric conditions. Moreover, disentangling the complexity of gene networks (and molecular pathways involved in these networks) could reveal new targets for pharmacological interventions or provide guidance to develop new personalized music-based therapies.

Recently, we demonstrated that musical stimuli have an important impact on the capillary blood transcriptomes of ACD patients and healthy individuals, providing insights into the systemic gene expression response to music. We reported that music stimulation in ACD patients compensates for the expression of genes and pathways dysregulated due to cognitive impairment. Elaborating on this groundwork and given the known connection between the oral cavity and the brain, namely oral-brain axis, we aimed for the first time to investigate gene expression changes elicited by music in saliva samples in healthy donors and ACD patients. For this purpose, we have followed the same experimental design that has already been successfully used in our previous study on capillary blood samples, but this time employing a specific saliva collection device and a hybridization-based and PCR-free *n*Counter assay from NanoString. Currently, this is the most appropriate technology to deal with transcriptomes isolated from saliva samples, which are usually enriched with abundant genetic material from microbial species and poor in terms of quantity/quality.

Overall, the results suggest that music significantly impacts on the salivary transcriptomes of patients and controls; with this impact being higher than in the capillary blood transcriptomes of donors (as evidenced by the number of DEGs captured, both non-adjusted and adjusted; **Table S2**). This finding highlights, for the first time, the relevance of the host transcriptome response to music in saliva. Additionally, the results indicate that the relative impact of music on individual genes may vary considerably between tissues, leading to specific genes being upregulated in one tissue but downregulated in another; this differential behavior between tissues is consistent with responses to other significant external stimuli in the host, such as an infection (28).

Three significant observations from the present study on the impact of music on salivary transcriptomes have already been reported for capillary blood transcriptomes (2), namely, music elicits: *i*) greater transcriptomic changes in ACD patients than in controls, *ii*) a transcriptomic response towards upregulation in ACD patients compared to healthy donors, and *iii*) music modifies the salivary expression of a few genes that are known to be altered in AD/MCI conditions. Despite the relatively low overlap existing between DEGs from saliva and blood capillary samples, *KDM6B* (Lysine Demethylase 6B) and *TIMP2* (TIMP Metallopeptidase Inhibitor 2) genes emerged as the only common DEGs (adjusted *P*-value < 0.05) in both tissues from ACD patients. *TIMP2* was downregulated in both tissues whereas *KDM6B* showed opposite regulation patterns between tissues after musical stimuli. Furthermore, these genes were also found significantly upregulated in MCI/AD patients compared to healthy controls (**Figure S1**). As reported in the literature, both *KDM6B* and *TIMP2* are required for a normal brain function, as alterations in these genes might lead to neurological conditions. *TIMP2* protein regulates extracellular matrix (ECM) remodeling and is particularly enriched in the hippocampus in comparison to other TIMP proteins (29). In the adult brain, *TIMP2* participates in neurogenesis, neuronal differentiation and in hippocampus-dependent memory (30, 31). Moreover, TIMP2 level disturbances were found across several neurodegenerative disorders such as AD (32-34). *KDM6B* cooperates with Tau in regulating synaptic plasticity and cognitive function (35) and is present in excitatory neurons. Deleting *KDM6B* in neurons led to impaired synaptic activity, resulting in learning and memory deficits in mice (35). Indeed, *KDM6B* has been recently reported as a risk gene for intellectual disability (36), highlighting its importance for an adequate brain activity.

In ACD patients, the *CXCL8* and *LGALS3* genes emerged as the most upregulated and downregulated genes in TP2, respectively. *CXCL8* gene encodes the pro-inflammatory IL-8 cytokine and is primarily expressed in neurons, astrocytes, and microglia in the nervous system. Chronic inflammation has been reported to be a pivotal factor in AD development (37), even from an early stage of the disease progression (MCI) and potentially preceding clinical symptoms. Significantly higher levels of CXCL8 have been reported in cerebrospinal fluid (CSF), brain and plasma from AD patients in comparison to levels from healthy controls (38-41), suggesting a probable detrimental role of CXCL8 in AD. *CXCL8* has been negatively correlated with cognitive scores from AD patients (38), and positively correlated with CSF Aβ levels (40). However, a neuro-protective role of CXCL8 in AD has also been suggested. While stimulation with amyloid beta (Aβ) triggers *CXCL8* production, it exhibits neuroprotective effects against Aβ-induced toxicity, possibly through a *CXCL8*-induced intracellular signaling and the production of neurotrophic factors, such as brain-derived neurotrophic factor (39). Thus, it is tempting to interpret that the upregulation of the *CXCL8* gene observed in saliva might represent a neuroprotective response triggered by music in the brains of ACD patients. Nevertheless, the specific role of *CXCL8* in AD pathogenesis is still unclear, and the disparity in results reporting both neuroprotective and detrimental effects may reflect issues related to e.g. different experimental conditions and tissues.

The main pathological characteristics of AD include the formation of Aβ plaques and neurofibrillary tangles, neuronal loss, inflammation, oxidative stress, and microglial activation. Galectin-3, encoded by *LGALS3* gene, has been extensively associated with the activation of microglial cells around Aβ plaques in AD, indicating a major involvement in disease pathogenesis (42-44). Molecular signatures of microglial activation in AD and ageing have been described *LGALS3* as one of the most upregulated genes (44, 45). Although there are no data on the Galectin-3 measurements in microglia from MCI patients, elevated galectin levels have also been reported in the serum of both AD and MCI patients (46-49), suggesting a role of Galectin-3 in the disease and/or a risk factor for disease development. These pieces of evidence support the usefulness of Galectin-3 as a biomarker for AD, and its inhibition could have significant therapeutic benefits (42). Therefore, the under-expression of *LGALS3* might suggest a beneficial effect of music in ACD by compensating for the pathological effect of *LGALS3* over-expression due to the disease condition.

The top DEG in healthy controls, the Thimet Oligopeptidase (*THOP1*) gene, showed a significantly lower expression after the musical stimuli. *THOP1* is responsible for encoding a metallopeptidase, which participates in the metabolism of different neuropeptides expressed in neurons and glial cells, and plays a role in the brain neuropeptide degradation (50). Dysregulation of *THOP1* has been associated with an unbalance in dopamine and serotonin turnover (51) and it is widely understood that listening to music can influence both dopaminergic and serotoninergic pathways. Thus, under-expression observed in the *THOP1* gene may be indicative of a music-mediated regulation of these neurotransmission systems. In addition, some studies have reported an over-regulation of *THOP1* in brain and CSF of AD patients (52, 53), indicating a potential association with Aβ-mediated toxicity. However, this over-expression of *THOP1* in AD patients has been attributed to a protective response against Aβ toxicity (54).

Pathways analysis of DEGs suggests an influence of musical stimuli on unsaturated fatty acid metabolism in ACD patients. The brain, predominantly composed of lipids, necessitates proper lipid homeostasis for a normal brain function and development. Within the brain, unsaturated fatty acids, particularly polyunsaturated fatty acids (PUFAs) govern critical processes such as cell survival, neurogenesis, brain inflammation and synaptic function (55). With normal ageing, there is a decline in cholesterol and PUFAs levels in lipid rafts, affecting cell-cell communication, signal transduction, and synaptic plasticity. However, this reduction is significantly more prominent in AD and other neurodegenerative diseases, leading to dysregulation in unsaturated fatty acid metabolism, increased APP processing, and rapid formation of Aβ aggregates (56, 57). Reduced levels of unsaturated fatty acids have been detected in the brain and plasma of AD patients (57, 58). Interestingly, various therapeutic approaches targeting lipid metabolism are being considered in the context of AD (59); our finding indicating a role of music in lipid homeostasis deserves further exploration.

Co-expression modules analysis pointed out to a high impact of music on the salivary transcriptomes, higher in ACD than in healthy controls (both in correlation values and lower *P*-values). Overall, the molecular response to music was characterized by a stronger involvement of both adaptative and innate immune systems in healthy controls than in ACD patients. Specifically, the over-regulated *GZMA* module was found to be engaged in several T-cell adaptative related responses whereas the under-regulated *SERPINA1* module participates in different innate processes, such as IL-1 and TLR signaling. Interactions between the immune and nervous systems are bidirectional, with each directly influencing the behavior of the other. Furthermore, both the adaptive and innate immune systems play complex and dynamic roles in learning and memory, brain function, and neurostimulation and collaborate closely to preserve immune homeostasis (60).

Dysregulation of these immune components can have significant repercussions on brain function and development, underscoring the importance of proper immune regulation in maintaining neurological health. For instance, microglia are macrophage-like innate resident immune cells of the central nervous system (CNS) with essential functions in the brain, ranging from immune surveillance and response to synaptic pruning and neuroprotection (61-63). Microglia interact with neurons modulating synaptic transmissions and, therefore, directly influencing synaptic plasticity and neuronal excitability (64, 65). Dysregulation of microglial activity has been implicated in neurological and neurodegenerative disorders, such as AD (66). Another innate immune component, the TLR family, is expressed in microglia, astrocytes and oligodendrocytes, but neurons also express TLRs, regulating proliferation, differentiation, outgrowth and neuron survival (67). TLRs cascades are involved in different brain-related functions contributing to neurogenesis, modulation of CNS plasticity and learning (68, 69). Disturbances in TLR signaling might have either a negative or positive impact on nervous system homeostasis. Studies also provide evidence of the important role of cytokines considered pro-inflammatory, such as the innate immunity mediators IL-1 and TNF, for normal synaptic function (70, 71). Similarly, adaptative immunity T-cells have showed to be important for normal brain functioning with a beneficial role in cognition and behavior (72-74). A decreased number or dysfunction of T-cells may contribute to the etiology of different neurological disorders like autism or AD.

Notably, *CD59* gene emerged as both one of the top downregulated DEGs and the hub gene of the under-regulated module in ACD patients. CD59 is a glycoprotein that plays a crucial role in regulating the complement system by preventing the formation of the Membrane Attack Complex (MAC). In the brain, complement proteins regulate neurodevelopment, neural migration, proliferation and synaptic pruning. Downregulation of *CD59* (75) and upregulation of the complement system have been reported in several studies involving AD mouse models and brain tissue from AD patients (76-78). However, it is still unclear if the changes in complement activity observed in AD are harmful or beneficial, as studies report both neuroprotective and neurodegenerative roles (77, 79-82).

Therefore, the biological interpretation of the expression changes induced by music is complex. The two downregulated modules in ACD patients and controls (*CD59* and *SERPINA1*) showed some functional commonalities, with intracellular transport and autophagic pathways emerging as common processes associated with the musical stimuli in both modules. Autophagy promotes the clearance of misfolded proteins and pathological aggregates, and in the brain, helps to maintain neuronal cellular morphology and physiological activities for a proper CNS function, prevents cellular toxicity and plays a crucial role in synaptic plasticity (83). In fact, deficient autophagic machinery is one of the most relevant hallmarks found in neurodegenerative diseases like AD (84, 85). The effect of music on the downregulation of modules involved in autophagy could be related to a local response, as these pathways are crucial for the homeostasis of most, if not all, tissues. However, this issue deserves further investigation in future studies due to the significant role of autophagy in neurodegenerative processes.

After our initial and recent attempt to investigate the impact of music on neurodegenerative diseases, the present follow-up study is pioneering in revealing several aspects: *i*) It demonstrates that music has the ability to influence the expression patterns captured from saliva donors, *ii*) It establishes parallels between gene expression observed in saliva and blood; *iii*) It highlights that music has a stronger impact on the transcriptome of ACD patients compared to healthy individuals, as e.g. measured by the number of DEGs altered by music, and *iv*) It reveals that music overall triggers the upregulation of gene expression in patients compared to healthy controls.

Among the limitations of the present study, we echo those already discussed in our previous study on sensogenomics22 experimental concerts carried out on capillary blood samples (2). Additionally, we acknowledge the challenge posed by the analysis of saliva in RNAseq analysis. Fortunately, the methodology employed in the present study, although it evaluates a lower number of genes for expression, offers the advantage of being a gold standard for gene expression studies. Therefore, the high quality of the gene expression results provided by NanoString compensates somewhat for the limitation of analyzing fewer genes.

The present study has demonstrated the power of short-duration musical stimuli in modifying the salivary transcriptome of ACD patients and healthy donors. The impact of music on saliva tissue is comparable to, or even greater than, that observed in blood with a more pronounced effect seen in patients than in healthy controls. Of note is the discovery that music influences the expression of genes and modules commonly altered in neurodegenerative diseases, a finding that may help to elucidate the known beneficial effects of music as reported by specialists in neuroscience and cognitive sciences. Further efforts to validate these findings in larger cohorts and other disease scenarios and to explore the impact of music not only on the gene expression level but also on other ‘-omic’ layers are warranted.

## Ethics approval and Consent to participate

Written informed consent was obtained from all the participants in the present study. The Ethics Committee of Xunta de Galicia approved the present project (Registration code: 2020/021), and the study was conducted in accordance with guidelines of the Helsinki Declaration.

## Consent for publication

All participants have given permission for publication of the project’s findings.

## Availability of data and materials

The authors confirm that data supporting the findings of this study are available in Gene Expression Omnibus – NCBI (GEO; https://www.ncbi.nlm.nih.gov/geo/) under accession number ###.

## Author’s contribution

AS, FM-T and LN conceived and coordinated the study; AS and AGC analyzed the data and wrote the first draft; AC-M, AD-U, AS, CM-C, CR-T, FC-V, FM-T, IF, IR-C, JM-L, JP-S, LC-M, LR, M-JC, NEZM, NM, SP, SR-V, SR-V, SV-L, XB, contributed to logistics and sample and data collection. All the authors revised and approved the final version of the text.

## Acknowledgements

We would like to kindly acknowledge all participants in the present study, the AGADEA association of Alzheimer disease patients, and the musicians of SANARTE, who kindly agreed to participate in this project. We would also like to acknowledge the Real Filharmonía de Galicia (www.rfgalicia.org; and Sabela García Fonte in particular), and the Auditorio de Galicia for their support. This study received support from GAIN (IN607B 2020/08 [A.S.] and IIN607A2021/05 [F.M-T]).

## Legend to the supplementary figures and tables

**Table S1.** Differential expression analysis results after comparing TP1 and TP2 in the age-related cognitive disorders (ADC) and control donor’s cohort.

**Table S2.** Proportion of DEGs (non-adjusted and adjusted) detected in capillary blood and saliva samples calculated using a) all detected genes and b) all common detected genes between both tissues. HC: Heathy controls; ACD: age-related cognitive disorders.

**Table S3.** Co-expression modules detected in ADC and control subjects.

**Table S4.** Gene Ontology (GO) and Reactome pathways significantly over-represented in the co-expression modules detected in ACD and control subjects.

**Figure S1.**
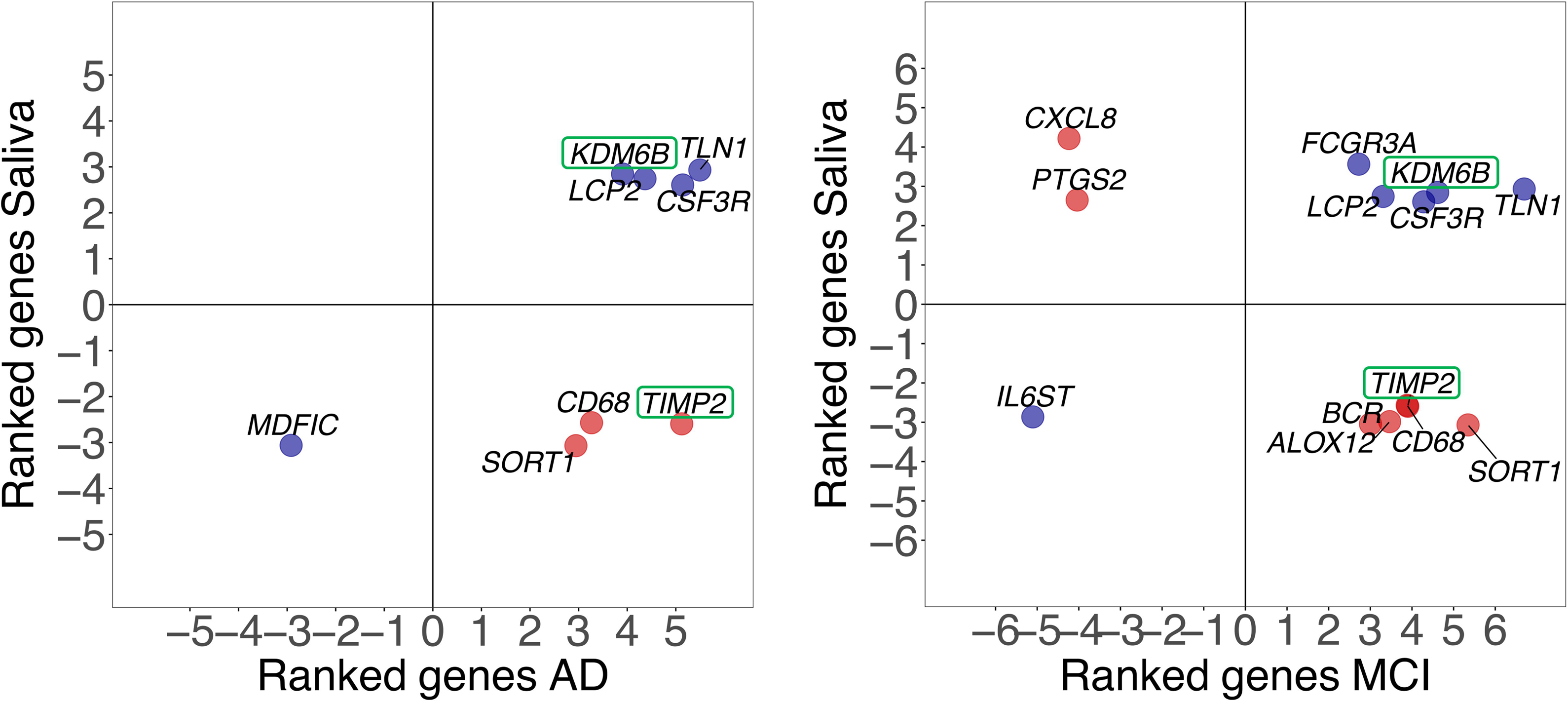
Correlation between gene expression changes observed in MCI/AD patients *vs.* healthy controls and saliva samples after musical stimulation in ACD patients. Blue dots indicate positive correlation between DEGs in the contrasts, while red dots indicate negative correlation. Labels of DEGs (adjusted *P*-value < 0.05) appearing in both MCI/AD and saliva samples are displayed. Ranked values refer to the value of the test statistic for a gene obtained from *DESEq2* for saliva samples and moderate t-statistics values obtained from *limma* for MCI/AD samples.

**Figure S2.**
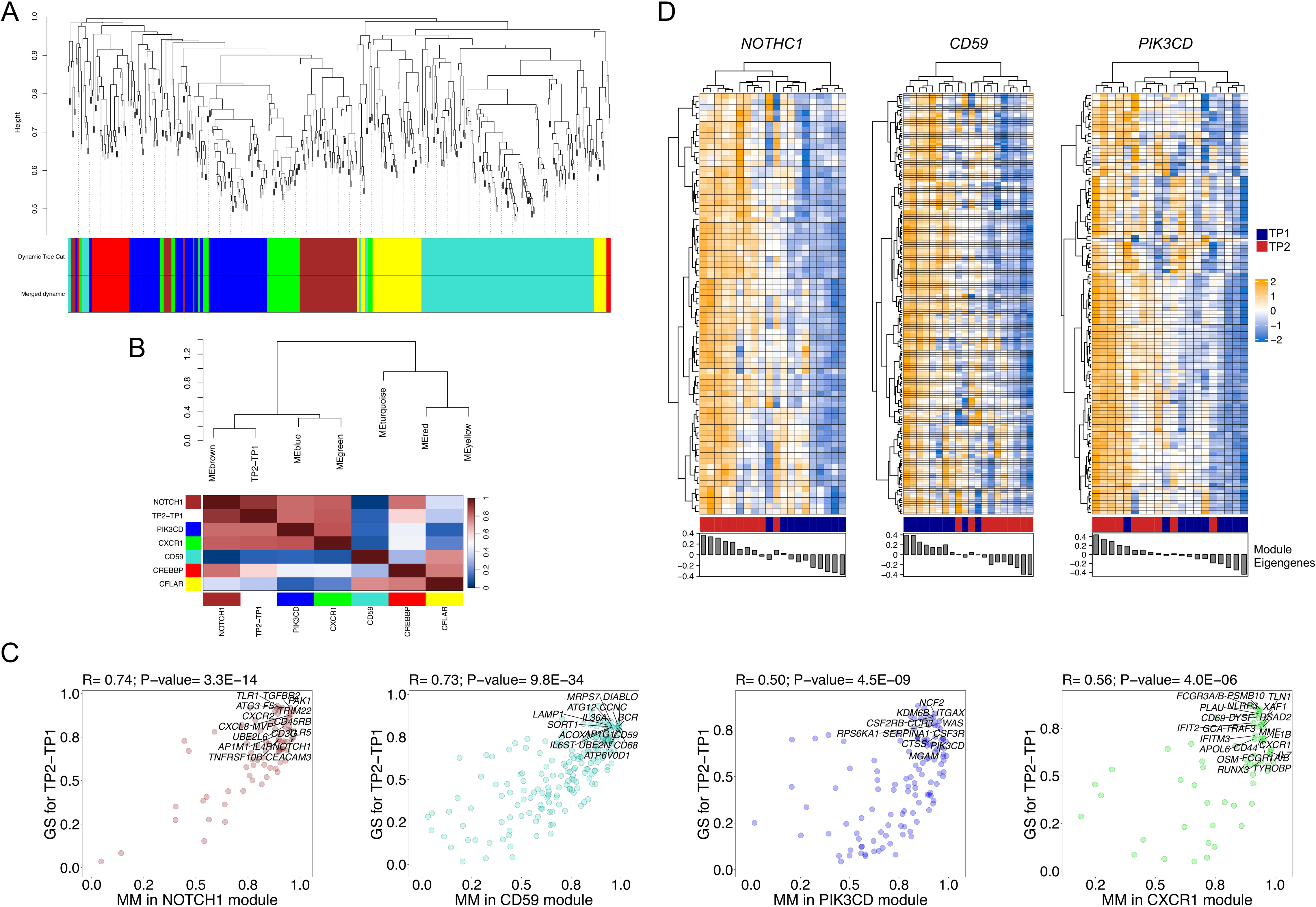
A) Clustering dendrogram of genes and co-expression modules detected in ACD patients represented by different colors. B) Eigengene dendrogram and heatmap for ACD patients displaying relationships among the modules and the trait of interest (musical stimuli; TP2 *vs.* TP1) in ACD patients. C) Module membership (MM) and phenotype (musical stimuli) correlation (GS) for genes from the statistically significant correlated modules in ACD patients. D) Gene expression heatmap and cluster analysis obtained from the genes included in the significant modules with significant pathways associated in ACD patients.

**Figure S3.**
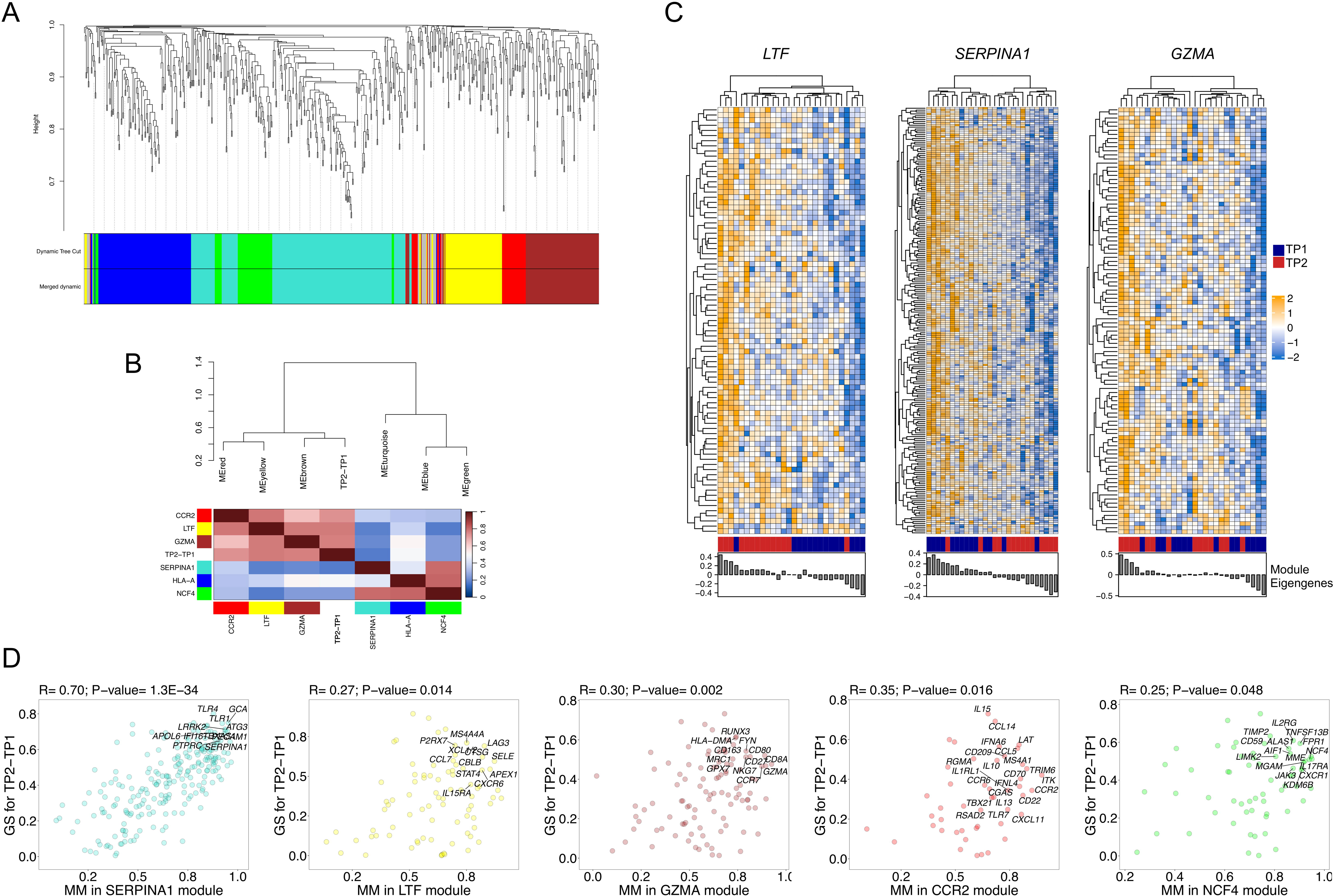
A) Clustering dendrogram of genes and co-expression modules detected in control donors represented by different colors. B) Eigengene dendrogram and heatmap for control donors displaying relationships among the modules and the trait of interest (musical stimuli; TP2 *vs.* TP1) in control donors. C) Module membership (MM) and phenotype (musical stimuli) correlation (GS) for genes from the statistically significant correlated modules in control donors. D) Gene expression heatmap and cluster analysis obtained from the genes included in the significant modules with significant pathways associated in control donors.

